# Methods for quantitative susceptibility and R2* mapping in whole post-mortem brains at 7T

**DOI:** 10.1101/2020.05.07.082479

**Authors:** Chaoyue Wang, Sean Foxley, Olaf Ansorge, Sarah Bangerter-Christensen, Mark Chiew, Anna Leonte, Ricarda AL Menke, Jeroen Mollink, Menuka Pallebage-Gamarallage, Martin R Turner, Karla L Miller, Benjamin C. Tendler

## Abstract

Susceptibility weighted magnetic resonance imaging (MRI) is sensitive to the local concentration of iron and myelin. Here, we describe a robust image processing pipeline for quantitative susceptibility mapping (QSM) and R2* mapping of fixed post-mortem, whole-brain data. Using this pipeline, we compare the resulting quantitative maps in brains from patients with amyotrophic lateral sclerosis (ALS) and controls, with validation against iron and myelin histology.

Twelve post-mortem brains were scanned with a multi-echo gradient echo sequence at 7T, from which susceptibility and R2* maps were generated. Semi-quantitative histological analysis for ferritin (the principal iron storage protein) and myelin proteolipid protein was performed in the primary motor, anterior cingulate and visual cortices.

Magnetic susceptibility and R2* values in primary motor cortex were higher in ALS compared to control brains. Magnetic susceptibility and R2* showed positive correlations with both myelin and ferritin estimates from histology. Four out of nine ALS brains exhibited clearly visible hyperintense susceptibility and R2* values in the primary motor cortex.

Our results demonstrate the potential for MRI-histology studies in whole, fixed post-mortem brains to investigate the biophysical source of susceptibility weighted MRI signals in neurodegenerative diseases like ALS.

## 1 Introduction

Magnetic resonance imaging (MRI) is able to provide a range of imaging contrasts sensitive to different properties of the tissue environment, and is thus a promising tool to study neurodegenerative diseases. However, even quantitative MRI methods are not intrinsically biologically specific, and interpretation of results requires comparisons against a more specific reference such as histology. In this study, we consider the use of susceptibility weighted MRI measures that are purported markers of iron and myelin in the context of a specific neurodegenerative disease, amyotrophic lateral sclerosis (ALS). In particular, we present methods for quantifying magnetic susceptibility and R2* in whole, fixed post-mortem brains, with direct comparisons to histology in the same tissue.

Susceptibility weighted MR data acquired using a gradient echo (GRE) sequence can provide several related but distinct quantitative measures. Magnitude images from GRE acquisitions at multiple echo times can be used to estimate maps of the effective transverse relaxation rate (R2*=1/T2*). R2* relates to compartmentalised “inclusions” with a susceptibility offset compared to the surrounding tissue (e.g. iron deposition) which broaden the distribution of frequencies within a voxel [1, 2]. R2* mapping benefits from relatively simple and robust processing. The GRE signal phase is driven by a (weighted) average magnetic susceptibility in a voxel, which relates to the relative size of each susceptibility-shifted compartment [3, 4]. Quantitative susceptibility mapping (QSM) employs sophisticated postprocessing on raw phase images to provide a quantitative estimation of the bulk susceptibility in an imaging voxel [5-8].

Susceptibility weighted MRI is sensitive to the para- and diamagnetic properties of tissue [9]. This sensitivity forms the foundation of contrast in images derived from both QSM and R2* mapping [5, 10-12]. In the brain, these maps have been shown to relate to the concentration of iron [13, 14] and myelin [15, 16] in tissue, which have para- and diamagnetic properties respectively relative to water. A fundamental difference between QSM and R2* mapping is that para- and diamagnetic inclusions have opposite effects on magnetic susceptibility (χ) estimates obtained from QSM, but the same effect on R2* rate. In neurodegenerative diseases, changes in tissue myelin and iron content may co-occur. As myelin and iron have opposing susceptibility signs (relative to water), the combination of QSM and R2* mapping has the potential to shed light on the overall pathology, provided their histopathological correlates can be clearly defined. For example, demyelination and accumulation of iron both lead to a positive shift in χ, but an opposing effect on R2* (decrease in R2* due to demyelination and increase in R2* due to iron deposition).

To date, several studies have reported the correlation of R2* and susceptibility with brain iron and myelin content using either whole, unfixed post-mortem brains (in situ) or small, fixed brain tissue samples [13-15, 17-20]; however, no studies have applied QSM in whole, fixed post-mortem brains. Whole, fixed brains present additional challenges including the presence of air bubbles, which induce rapid focal field variations, and the influence of brain shape on the macroscopic magnetic field homogeneity, which can make shimming difficult. To directly compare whole-brain QSM to histology in the same tissue, we therefore need to develop bespoke imaging approaches and processing pipelines for whole, fixed post-mortem brains.

Here, we demonstrate this approach in an exemplar neurodegenerative disease. ALS is a neurodegenerative disease of the motor system, characterized by rapidly progressive and irreversible loss of motor function, ultimately leading to death [21]. Similar to other neurodegenerative diseases, ALS pathology is believed to follow a standard progression within the cerebral cortex that starts in the primary motor cortex (M1) [22, 23] and gradually progresses into non-motor cortices [22, 23]. In end-stage disease (corresponding to post-mortem brains), M1 would be expected to exhibit substantial pathology, anterior cingulate cortex (ACC) to exhibit intermediate pathology, and secondary visual cortex (V2) to be largely unaffected [23]. Therefore, development of MRI biomarkers related to these regions would be helpful for studying disease mechanisms; for tracking disease progression and response to therapeutics; and potentially for early diagnosis. A study comparing R2* mapping with iron histology in a single ALS brain demonstrated elevated R2* in the deep layers of M1 that corresponded to iron accumulation [24]. Recent in vivo QSM studies of ALS have shown elevated susceptibility in the motor cortex and a range of subcortical nuclei [25-27], hypothesised to be driven by an increased concentration of iron.

We present a pipeline for comparing χ and R2* to semi-quantitative histology in whole, fixed post-mortem brains. Our motivation for scanning whole brains, as opposed to small tissue samples, is the ability to study the relationship between MRI and histopathology across different brain regions that are affected at different disease stages [28]. Susceptibility weighted MR images were acquired using a multi-echo gradient echo (GRE) acquisition in whole, fixed post-mortem brains from nine ALS patients and three control subjects. These data were subsequently processed using a custom analysis pipeline to generate quantitative susceptibility and R2* maps. Brains were then sectioned and stained using immunohistochemistry targeting myelin proteolipid protein (PLP) and ferritin in three brain regions with a different pathological stage profile (M1, ACC and V2). Our results provide proof-of-principle for using MRI-histology comparisons to study the histopathological correlates of neurodegeneration at the whole brain level.

## 2 Materials and Methods

### 2.1 Post-mortem brain samples

Post-mortem brains were obtained from the Oxford Brain Bank, University of Oxford, under its generic Research Ethics Committee approval (15/SC/0639). In total, twelve whole, post-mortem formalin fixed brains were scanned with our multi-echo GRE acquisition. The cohort consisted of nine donors with a clinical diagnosis of ALS, and three control donors with no known neurological disorder. Diagnosis was confirmed by a neuropathologist. Patient demographics, post-mortem information, and relevant tissue handline properties are listed in Table 1.

**Table 1.** Patient demographics and post-mortem information.

| Case# | Age | Sex | PM delay<br>(days) | Fixative | Time in<br>fixative before<br>scan (days) | Neuropathological<br>diagnosis |
| --- | --- | --- | --- | --- | --- | --- |
| 1 | 66 | M | 7 | 10% Formalin | 137 | ALS |
| 2 | 65 | M | 4 | 10% Formalin | 94 | ALS |
| 3 | 76 | M | 3 | 10% Formalin | 158 | ALS |
| 4 | 68 | M | 1 | 10% NBF | 35 | ALS |
| 5 | 54 | M | N/A | 10% Formalin | 147 | ALS |
| 6 | 65 | M | N/A | 10% Formalin | 96 | ALS |
| 7 | 68 | M | 2 | 10% NBF | 87 | ALS |
| 8 | 71 | M | 3 | 10% NBF | 94 | ALS |
| 9 | 79 | F | 2 | 10% NBF | 139 | ALS |
| 10 | 61 | M | 3 | 10% NBF | 115 | Control |
| 11 | 74 | M | 3 | 10% NBF | 45 | Control |
| 12 | 61 | M | 3 | 10% NBF | 48 | Control |

*N/A = Not Available; NBF = Neutral Buffered Formalin*

### 2.2 Sample preparation and MRI data acquisition

Brains were imaged in a 7T whole-body human scanner (Siemens Healthcare, Erlangen, Germany) using a 1Tx/32Rx head coil and a multi-echo GRE sequence. To prepare each brain for scanning, the brain was first removed from storage in formalin. Excess formalin remaining within the ventricles was removed by mildly compressing the brain whilst being held with the cerebellum pointing downwards. Excess formalin on the brain surface was removed using paper towels. The brain was then placed in a plastic bag filled with a susceptibility-matched perfluorocarbon liquid (3M™ Fluorinert™ FC-3283) and gently compressed/rotated to fill the ventricles with Fluorinert and remove any excess air-bubbles. This process also removed any remaining formalin within the ventricles, which has a lower density than Fluorinert (and hence floats to the top of the bag). This process continued until no new air-bubbles/formalin were observed. The brain was subsequently transferred and sealed in a 3D printed brain-shaped container (Figure S1 in the Supplementary material), keeping the fourth ventricle pointing upwards to prevent loss of Fluorinert during the transition. The 3D printed shell was subsequently filled with Fluorinert and gently manipulated to remove any new air bubbles that may have formed during the transfer.

This brain-shaped container was then placed in an external brain holder with a spherical chamber (Figure S1 in the Supplementary material) that was again filled with Fluorinert. This exterior holder achieves two aims: first, ensuring that brains are consistently positioned in the main magnetic field; and second, providing a spherical overall shape that minimizes field distortions in the brain. Both of these aims will benefit reproducible and high-quality susceptibility weighted data.

Susceptibility weighted data were collected using a 3D multi-echo GRE sequence with following parameters: TEs = 2, 8.6, 15.2, 21.8, 28.4, 35 ms with monopolar readout and non-selective RF pulse, TR = 38 ms, flip angle = 15°, bandwidth = 650 Hz/pixel, and GRAPPA factor = 2. Cases #1-7 were scanned with voxel size = 0.5×0.5×1.2 mm^3^ and matrix size = 384×310×104, total scan time = 11 min. Cases #8-12 were scanned with voxel size = 0.5×0.5×0.6 mm^3^ and matrix size = 384×310×208, total scan time = 22 min. GRE acquisitions were repeated four times for all brains and raw, multi-channel k-space data were collected. To perform comparative analyses at the same voxel size, the raw k-space data from Cases #8-12 were subsequently truncated to downsample the data to match the voxel size of Cases #1-7. As part of a larger study, structural, relaxometry and diffusion data were also acquired [28].

### 2.3 QSM post-processing pipeline

#### 2.3.1. GRAPPA reconstruction and image registration

Due to the need to work with raw k-space data in order to optimise coil combination, GRAPPA reconstruction was performed using an off-line MATLAB implementation with a single kernel across all echoes. After GRAPPA reconstruction, image misalignments between different echoes and different repeats were observed, which were highly consistent between brains. This misalignment was hypothesised to be induced by *B*_0_ drift caused by scanner temperature changes over the entire experiment for each brain [29]. As the phase from 1^st^ echo was only utilized to remove coil-specific phase offsets in the coil combination step (described below), in order to minimalize data interpolation due to registration, linear image registrations were performed using FLIRT (FMRIB’s Linear Image Registration Tool) [30-32] to register the 1^st^ echo to 2^nd^ to 6^th^ echoes respectively. This was done separately for the 4 repeats.

#### 2.3.2 Coil combination

The images from a single RF channel were found in many spatial locations to exhibit open-ended fringe lines (non-physical phase singularities) in the region of highly focal signal loss. In addition, the time-independent phase offset, 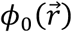 is unique for every channel and needs to be accounted for when combining multi-channel data.

The phase offsets for each channel was removed by subtracting the phase of the co-registered 1^st^ echo *ϕ*(*TE*_1_) (*TE*_1_ = 2 *ms*) from *ϕ*(*TE*_2_) to *ϕ*(*TE*_6_), producing five phase images Δ*ϕ*(*TE*′) for each channel with equivalent echo times *TE*′ of 6.6, 13.2, 19.8, 26.4 and 33 ms and no time-independent phase offset. For each equivalent echo time, a coil-combined phase image was generated using a complex sum over all the coils [33]. Magnitude images were combined using a simple “sum-of-squares” method to produce the magnitude data of all 6 echoes.

#### 2.3.3 Phase unwrapping and field map estimation

As image misalignments (likely due to *B*_0_ drift caused by scanner temperature changes) were observed, another registration step was performed to align all coil-combined phase images Δ*ϕ*(*TE*′) from the 4 repeats using FLIRT [30-32], which used one echo from one repeat as a reference and registered all other echoes from the 4 repeats to the reference echo.

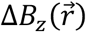 was estimated using a voxel-wise non-linear fit of the complex data, 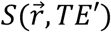, over the co-registered coil-combined echoes [34, 35] by minimizing:

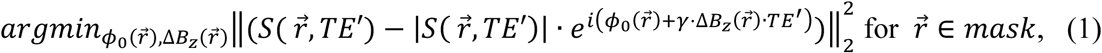

where *γ* is the gyromagnetic ratio and 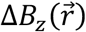 is the z-component of the magnetic field perturbation. The cost function is evaluated only for voxels in a mask that was generated by thresholding the magnitude image from the first echo. Assuming linear phase evolution, *ϕ*_0_ will have been removed by our coil combination. We include 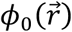 in the 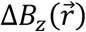 fitting process as an indication of the goodness of the fit. Small remaining estimates of *ϕ*_0_ may correspond to a slightly non-linear phase evolution due to tissue microstructure. However, any large, focal estimates of *ϕ*_0_ would be indicative of noisy voxels with unreliable phase, which should not be included in the later steps of QSM processing. Fitting was performed using the *lsqnonlin* function in MATLAB (R2016a).

Fitting was initialized with estimates of 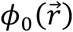 and 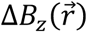 obtained from the first two echoes of our coil combined data. To achieve this, a phase difference map between the first two echoes was generated:

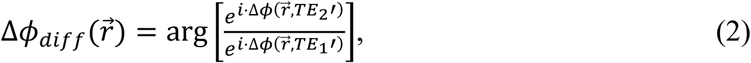

and spatially unwrapped using PRELUDE (Phase Region Expanding Labeller for Unwrapping Discrete Estimates) [36]. The phase map at the first echo was also spatially unwrapped and our initializations were estimated as follows:

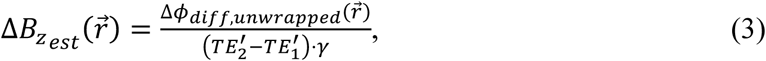

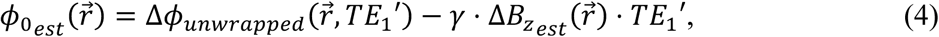

This initialization offers some advantages. Firstly, without initialization of the fitting, phase wraps may remain in the estimated 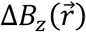 map due to rapid temporal phase evolution in tissue that exceeds the Nyquist limit. By providing an initialization consisting of a 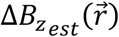 map on the order of the true 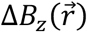, we can produce wrap-free 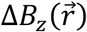 maps that are estimated from all the echoes in our offset-corrected phase data. Secondly, spatial phase unwrapping is only performed on the first echo and the phase difference between the first two echoes to generate 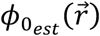 and 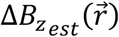. These images will contain few phase wraps compared to the phase images acquired at higher echo times, enabling robust unwrapping performance and initialization estimates with minimal artefacts.

As shown in Figure S2 in the Supplementary material, voxels with very large and focal 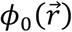 values occurred at the edges of the brain and were found to be due to the presence of vessels and air bubbles. These voxels are sources of artefacts of no interest in QSM. Therefore, the brain mask was refined by thresholding the gradient of this 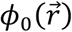 map to remove these focal field outliers.

When comparing the predicted phase evolution from *TE*′_1_ to the observed phase evolution in later echoes, we consistently observed a linear phase gradient along the readout direction (Figure 1, similar to the effect found by Tendler et al. [37]), which was hypothesised due to eddy currents or an accumulative mistiming of the centre of readout gradient (resulting a shift of k-space centre). These phase gradients would lead to apparent gradients in field offset along the readout direction and need to be removed. These shifts were estimated by a 1D linear fit along the readout direction over the phase increments of Δ*ϕ*(*TE*′_2_) to Δ*ϕ*(*TE*′_5_) with reference to Δ*ϕ*(*TE*′_1_), and subsequently removed from Δ*ϕ*(*TE*′_2_) to Δ*ϕ*(*TE*′_5_).

**Figure 1.**
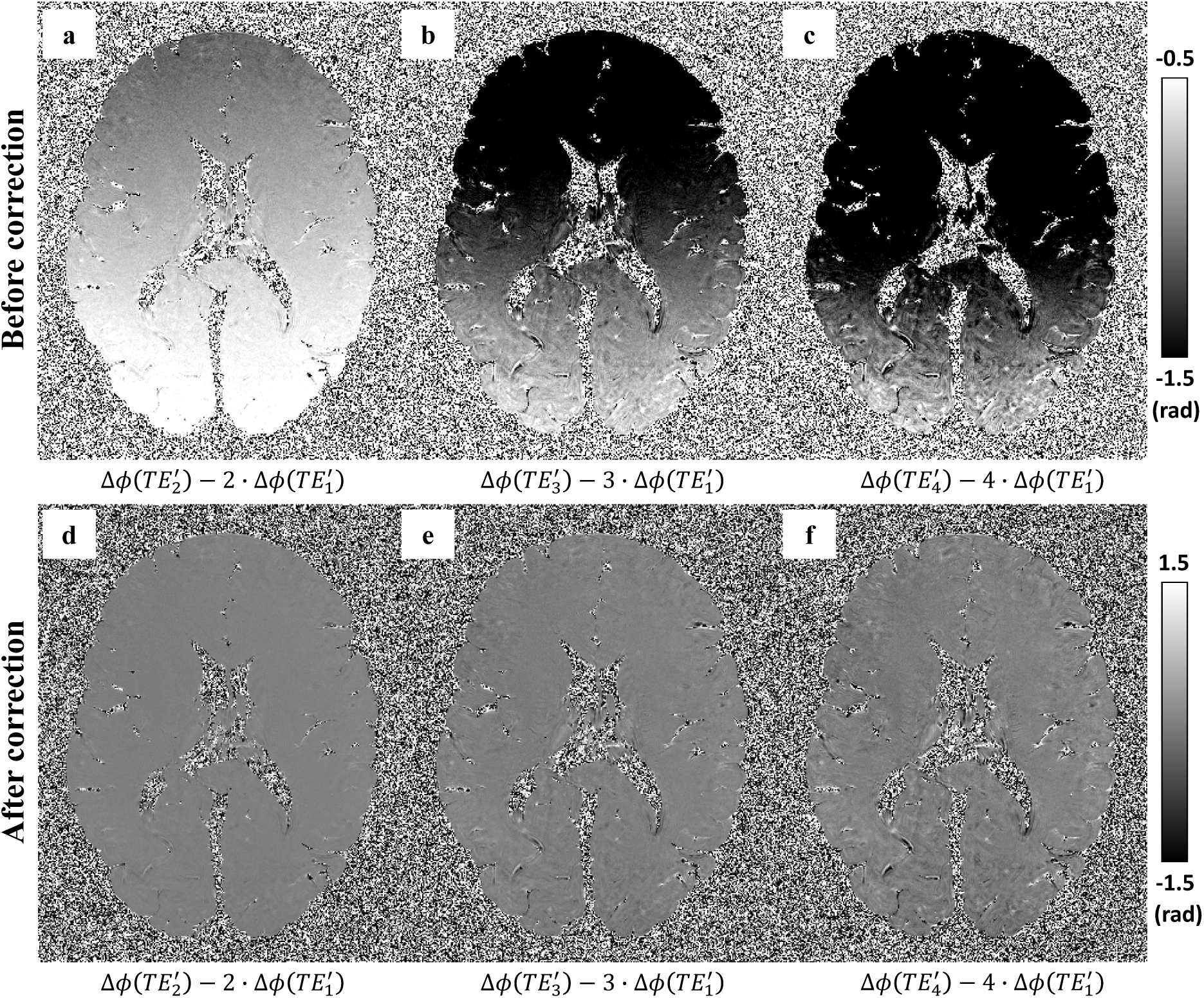
Difference between the observed phase and predicted phase based on the first effective echo *TE*′_1_ assuming linear phase evolution (Δ*ϕ*(*TE*_*n*_′) = *n* · Δ*ϕ*(*TE*_1_′)). If phase evolution was purely linear, these difference maps would be flat and close to zero. In practice, these phase difference maps consistently exhibit a linear phase gradient along the readout direction (a-c). A 1D linear phase was estimated and removed along the readout direction using the methodology described in [37], resulting in much flatter phase difference maps (d-f).

#### 2.3.4 Averaging, background field removal and dipole inversion

Four repeats of our GRE sequence were obtained per post-mortem brain sample (except Case #2, where only data from two repeats were available). Prior to each repeat, the scanner performs a whole-brain center frequency calibration, such that the phase over the whole brain should average to zero for each echo. Investigation of this mean temporal phase evolution over the four repeats instead revealed a non-zero, variable slope of phase evolution that consistently increased from the first to last repeat (quantified as the mean frequency offset of the 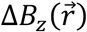 map, Figure 2). This is likely due to *B*_0_ drift induced by temperature drift in the scanner during acquisition [29]. These GRE data were obtained as part of a larger study [28] acquiring multi modal MRI data within each brain. Over the course of this acquisition, the gradient duty cycle ranges from an intensive 24-hour diffusion-weighted protocol to a moderate T1/T2 weighted protocol, making it likely that the scanner temperature was unstable during the GRE acquisition. Even with an accurate frequency calibration at the beginning of each GRE repeat, this could lead to frequency drift during each repeat.

**Figure 2.**
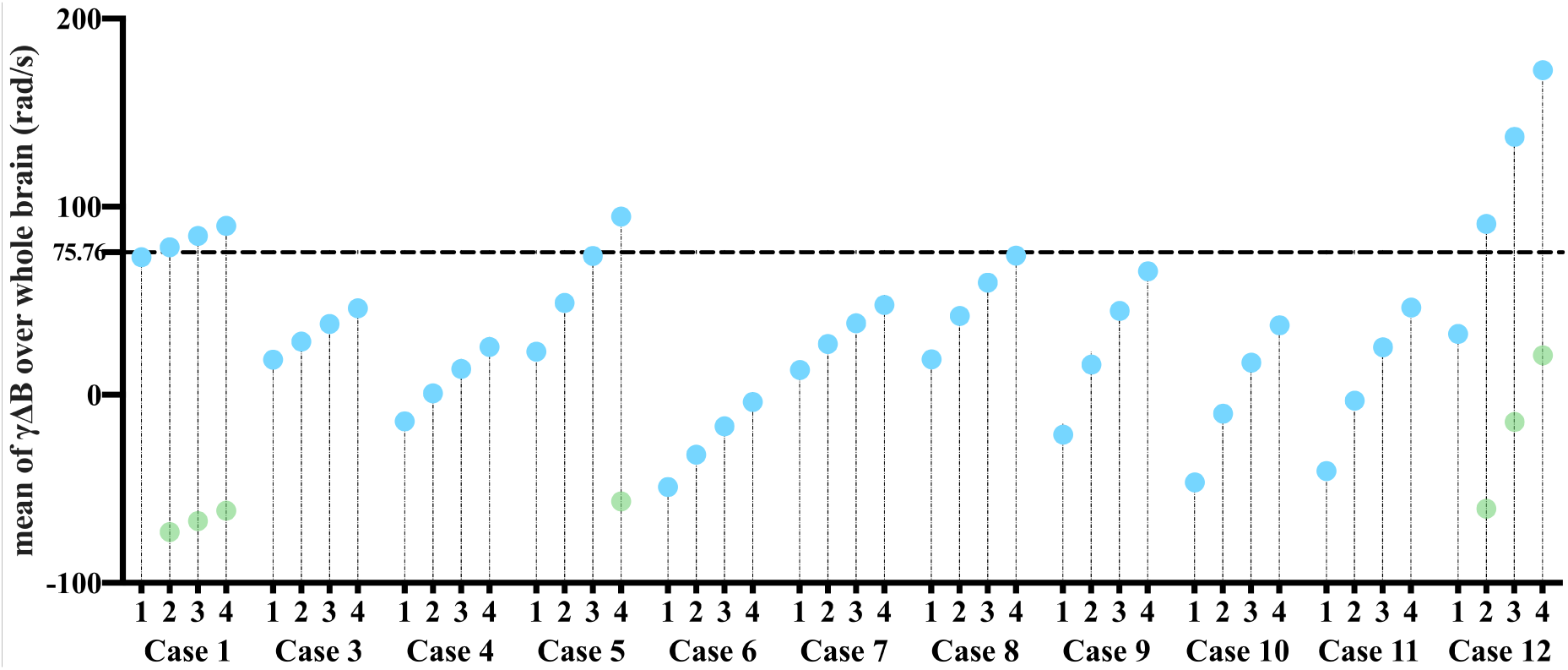
Mean values of the frequency evolution (Δ*f* = *γ*Δ*B*) over the whole brain for each of the 4 repeats (only two repeats were available for Case #2, which is not presented here). The dashed horizontal line corresponds to the Δ*f* value that accumulates phase of *π* with echo spacing of 6.6 ms 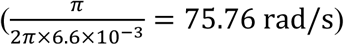, which leads to aliasing when Δ*f* is above this value (indicated by the light green dots). This was observed in Cases #1, #5 and #12.

Averaging the phase data from 4 repeats prior to fitting for 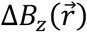, which would require a correction for this *B*_0_ drift to avoid phase cancellation. Instead, the estimated 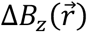 maps from 4 repeats were estimated and averaged to obtain the final 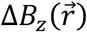 map. We found the latter approach to be more robust than the former to certain artefacts in the data. In particular, one brain showed evidence of Fluorinert leakage during the course of the scan, leading to local changes in 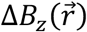 due to the difference in susceptibility between Fluorinert and air.

The experimental setup developed here differs from in vivo imaging in several important ways of relevance to QSM processing and algorithms that are optimized for in vivo imaging may not be appropriate post-mortem. Hence, an evaluation of the robustness of different algorithms for background field removal and dipole inversion in post-mortem data processing was performed. Three existing algorithms for background field removal were evaluated: variable kernel SHARP (v-SHARP) [38, 39], projection onto dipole fields (PDF) [40] and Laplacian boundary value (LBV) [41]. We also evaluated three algorithms for dipole inversion: thresholded k-space division (TKD) [42], morphology enabled dipole inversion (MEDI) [43] and streaking artifact reduction for QSM (STAR-QSM) [44]. Representative susceptibility maps from these comparisons are shown in Figure 3. Overall, we found that the combination of v-SHARP and STAR-QSM was the most robust for our given post-mortem datasets, yielding susceptibility maps which did not contain large scale inhomogeneities, had few steaking artefacts and did not appear over-smoothed.

**Figure 3.**
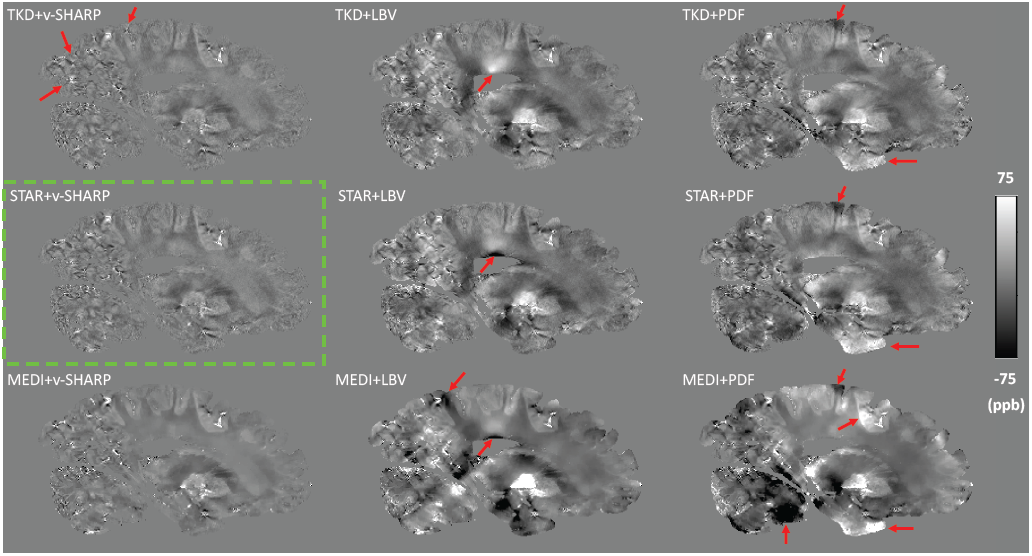
Susceptibility maps from combinations of three background field removal methods and three dipole inversion methods. Red arrows indicate streaking artefacts or remaining large scale inhomogeneities on other susceptibility maps compared to maps generated using STAR-QSM and v-SHARP. Susceptibility maps generated with MEDI generally showed over-smoothed contrast, for instance, there was less contrast within white matter regions but very sharp edges at the grey and white matter boundary.

Thus, background fields in the estimated 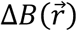 maps were removed using the v-SHARP algorithm [38, 39] with our refined mask, a maximal kernel size of 12mm and regularisation parameter of 0.02mm^-1^. Finally, the susceptibility (Δ*χ*) maps were generated using the STAR-QSM algorithm [44].

In QSM, the centre of k-space of the estimated susceptibility map is undefined [3], which makes it a measure of tissue’s relative susceptibility, an internal reference region is usually used, thus, the measured susceptibility values were referenced to the whole brain by setting the mean susceptibility value across the whole brain to zero.

### 2.4 R2* post-processing pipeline

This same multi-echo GRE scan was used to generate R2* estimates from the magnitude images. R2* maps were estimated using voxel-wise fitting of the magnitude images of all 6 echoes by minimizing:

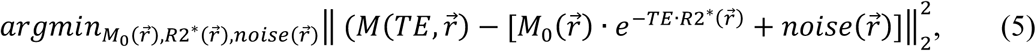

where 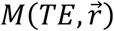 is the experimental magnitude data, 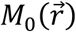 is the extrapolated signal at TE = 0 ms, and 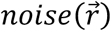 represents the noise floor.

As described above, *B*_0_ drift induced effects were observed and corrected during the processing of phase data. This is likely due to temperature change during scanning, which followed a long diffusion protocol with significantly higher gradient duty cycle than the GRE scans. Previous work has reported a linear dependence of T2* measurements on temperature [45]. To test for such effects in our data, the T2* maps from individual repeats of each brain were estimated and the mean T2* values over whole brain were calculated. Our results found a small change (0.41% ± 0.29%) in the mean T2* values from repeats #4 relative to repeat #1 over the 11 brains (change in ms for each brain and repeat shown in Figure S3 in the Supplementary material - only two repeats were available for Case #2, which was not included here). As the mean T2* value drifts over 4 repeats were small, no correction of the temperature effect was performed. The final R2* maps were generated using averaged magnitude images over the four repeats for each brain.

The processing pipeline for generating QSM and R2* maps is displayed in Figure 4.

**Figure 4.**
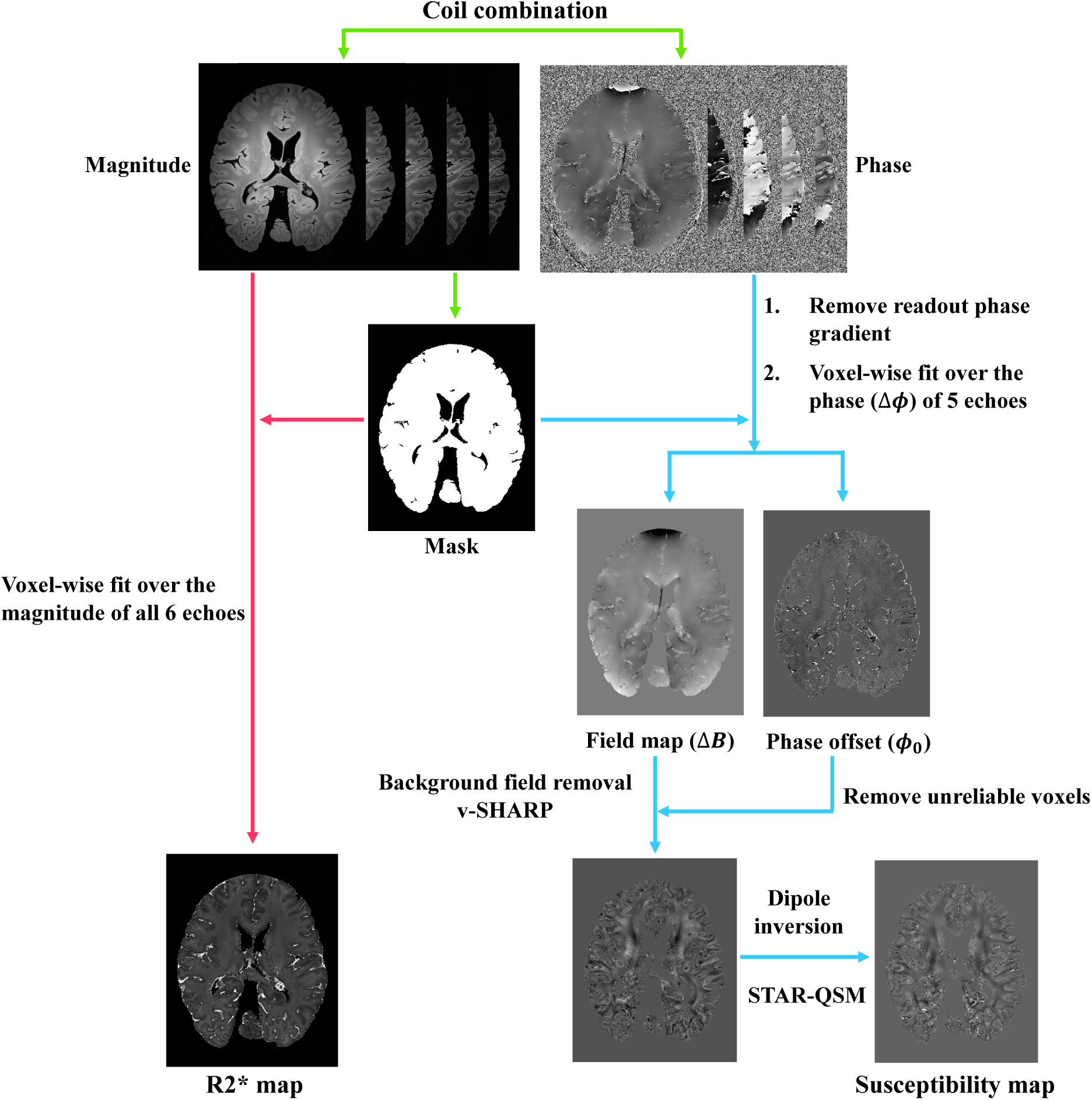
Illustration of the post-processing pipeline. Green arrows indicate the pre-processing steps including coil combination and mask generation, red arrows indicate the R2* mapping processing steps, and blue arrows indicate the QSM processing steps.

### 2.5 MR image analysis

Three regions of interest (ROIs) were analyzed (Figure 5) in the MRI data. These regions were chosen to represent different stages of ALS: M1 (Stage I), ACC (Stage II and III) and V2 (no pathology, as internal control). M1 was further separated into face, hand and leg areas. Masks of M1 regions were created in MNI space and subsequently registered into individual diffusion space and manually refined in each hemisphere by an experienced researcher familiar with M1 anatomy (R.M.) using the mean diffusivity maps from the diffusion acquisition as a reference. ROI masks were subsequently registered to the GRE data using FLIRT [30-32]. Masks of ACC and V2 were hand-drawn based on the location of the tissue block stained for histology. Briefly, the histological stained slides were carefully mapped to the corresponding location in the 3D structural images by visually comparing photographs of the coronal sampled brain slices to the MR image. Masks of ACC and V2 were manually drawn in the structural space to match the cortical region on the stained slides (described in Section 2.6), and finally registered to GRE data using FLIRT [30-32].

**Figure 5.**
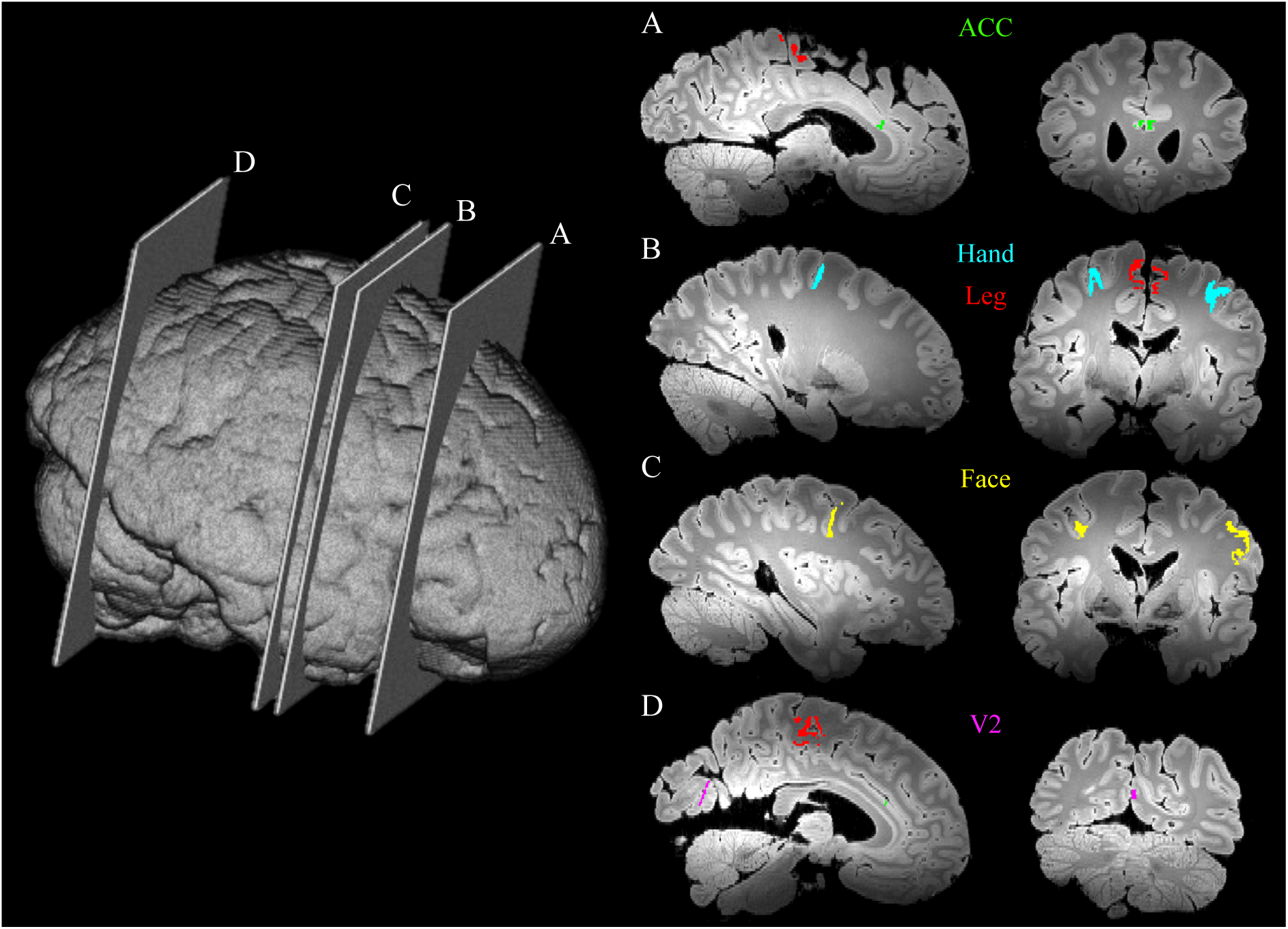
Illustration of slide sample locations across a whole brain from a single subject to highlight the ROIs used in our analysis. The yellow, blue, red, green and purple regions correspond to the M1 face area, M1 hand area, M1 leg area, anterior cingulate cortex (ACC) and secondary visual cortex (V2), respectively.

Due to a change in brain bank protocols, two types of fixative (10% neutral buffered formalin (NBF) and 10% formalin) were used for fixation of the brains. The effects of the two types of fixative on R2* and susceptibility values were analysed. As shown in Figure 6A and 6B, differences were found between two fixative groups in the mean R2* value over the whole brain (29.7% difference between the group means) and the M1 region (29.4% difference between the group means). Fixation effect (characterized as an indicator variable) was regressed out from mean R2* value over the whole brain. As the mean susceptibility value across the whole brain was set to zero for each brain, only the group difference in averaged susceptibility value in the M1 region between the two fixative groups was investigated (Figure 6C) and the group difference was found to be not significant (7.5% difference between the group means).

**Figure 6.**
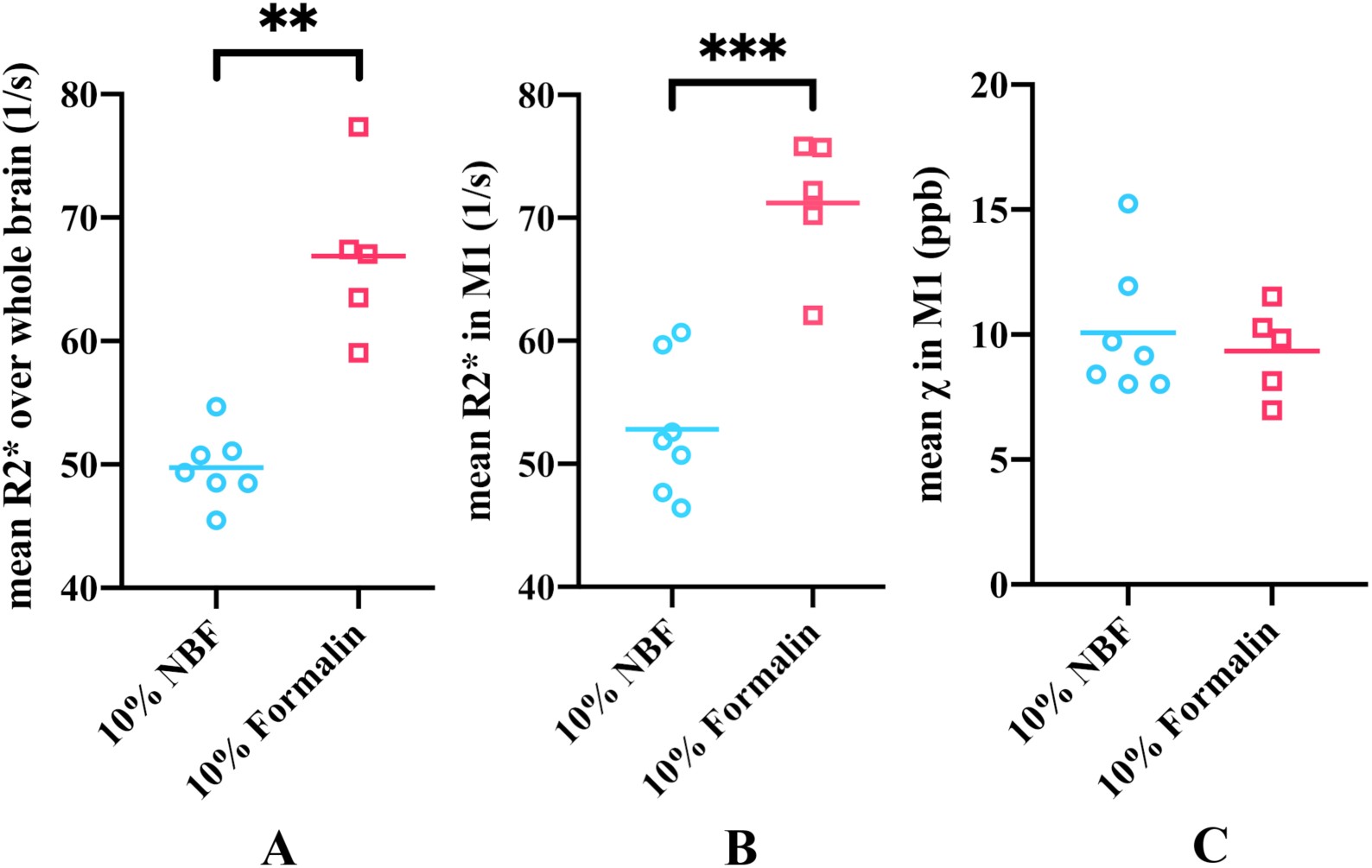
Comparison of the effects of the two types of fixative on susceptibility and R2* maps. Mean R2* values over the whole brain is plotted for the two types of fixative (A). Mean R2* (B) and susceptibility (C) values in the whole primary motor cortex (M1) region are plotted for the two types of fixative. Mean R2* values in the 10% neutral buffered formalin (NBF) group were significantly lower compared to the 10% formalin group over whole brain (p=0.0031, Hedges’s g=-3.287) and in M1 (p=0.0004, Hedges’s g=-3.059), whereas no significant difference was found for the mean susceptibility values in two fixative groups (p=0.58, Hedges’s g= 0.286) in M1. ** *p<0.01; *** p<0.001*

Effects of post-mortem delay and time in fixative before scanning on the susceptibility, R2* maps were also investigated: no significant associations were found.

Although care was taken to detect and remove the voxels corresponding to air bubbles as described above, some large positive or negative susceptibility values induced by remaining air bubbles still existed in the final susceptibility maps. An outlier detection process was performed on the susceptibility and R2* values within our anatomically defined ROIs prior to measuring the mean value. Outliers were defined as values more than three scaled median absolute deviations away from the median. These outliers were excluded from the ROI analysis. Following outlier rejection, mean susceptibility and R2* values within each ROI mask were calculated.

### 2.6 Histological analysis

Following MRI, the brains were systematically sampled and ROIs were extracted as described by Pallebage-Gamarallage et al. [28]. Briefly, the M1 leg region was represented on the medial surface of the paracentral lobule at the banks of the interhemispheric fissure. The M1 hand area (hand knob) was recognised by the inverted omega on the precentral gyrus at the anterior bank of the central sulcus. Similarly, the M1 face region was identified on the precentral gyrus, at the anterior bank of the central sulcus, lateral to the intersection of inferior frontal sulcus and precentral sulcus. Subsequently, the brains were sliced in a coronal plane for the ACC and V2 to be sampled. The ACC was extracted from the coronal slice represented with the striatum (caudate and putamen) and the anterior limb of the internal capsule rostral to the anterior commissure and posterior to the head of the caudate nucleus. V2 was sampled adjacent to the primary visual cortex (V1), identified by the stria of Gennari at the banks of the calcarine fissure. Sampled tissues were processed for paraffin embedding and 6 μm thick paraffin sections were cut and mounted on 75 × 26 mm slides for immunohistochemistry. Sections were deparaffinised in xylene, rehydrated in graded ethanol and water. Endogenous peroxidase activity was blocked by 3% H2O2 (in phosphate buffered saline) for 30 min prior to carrying out microwave antigen retrieval in citrate buffer. Sections were incubated with primary antibodies against ferritin (1:3000, Sigma F5012) and PLP (1:1000, Bio-Rad MCA839G) for 1 h. Proteins were visualised with DAKO EnVision Detection Systems where secondary antibody HRP (horseradish peroxidase) rabbit/mouse serum was applied for 1 h followed by 3,3’-diaminobenzidine (DAB) chromogen. Sections were counterstained with haematoxylin, dehydrated with graded ethanol, cleared in xylene and mounted with DPX mounting medium.

Whole slides were digitised at x20 objective magnification with Aperio ScanScope®AT Turbo (Leica Biosystems) high throughput slide scanner. Previously validated [28] preset Aperio Colour Deconvolution algorithm thresholds (version 9.1, Leica Biosystems) unique to each stain were applied to quantify the stained area fraction (SAF). The SAF is the ratio of positively stained area normalised by the total area within a region of interest (ROI). Cortical ROIs were defined by an experienced histologist based on histological and neuroanatomical landmarks on each slide (as described previously in Section 2.6). The annotations of cortical ROIs on histology slide are shown in Figure S4 in the Supplementary material. Representative digital histology and markup images of ferritin and PLP stains are shown in Figure S5 in Supplementary material. The SAF calculations were validated by a neuropathologist. Note that the measured SAF values have no units.

Ferritin staining was performed in the left hemisphere for M1 (leg and face areas, in two separate experiments), ACC and V2. In order to compare across regions in light of high cross-batch variability in ferritin staining, we conducted ferritin staining in two batches containing all samples, with each batch including one M1 region, ACC and V2. As a result of this cross-batch variability, the ferritin SAF calculated from two batches had substantial differences in the range of SAF values. To combine the ferritin results across experiments, SAFs were standardized within each batch (demean then divide by the standard deviation) prior to combined analysis of both batches. Thus, ferritin SAFs reported have zero mean, and therefore will include negative “fractions”. PLP staining was also performed in both hemispheres for M1 (leg, face and hand areas), ACC and V2.

No significant effects were found when comparing fixative type, post-mortem delay, or time in fixative before scanning for the histology data.

### 2.7 Statistical analysis

All statistical analyses were performed using SPSS version 25.0 (IBM, Armonk, NY). Pearson correlation analyses were used to assess (i) the relation between susceptibility and R2* values, (ii) the relation between ferritin and PLP, and (iii) the relation between pairs of MRI and histological stains.

Welch’s t-tests were used to compare (i) the group differences (ALS vs controls) in mean susceptibility and R2* in the M1 and (ii) the group differences (ALS vs controls) in ferritin and PLP stains in available M1 areas. Hedges’s g was calculated for each t-test as effect size, which has a similar interpretation to Cohen’s d but is generally considered more accurate for small sample sizes.

All p-values presented are for two-tailed tests. A p-value less than 0.05 was considered significant.

## 3 Results

### 3.1 Susceptibility and R2* maps

Figure 7 depicts the four ALS cases that exhibited visually apparent hyperintensities in susceptibility and R2* within the cortical grey matter of M1 compared to the surrounding tissue, including the adjacent primary sensory cortex. For example, this is clearly visible around the ‘hand knob’ in Case #6 and #9, but also extending towards face and leg areas. The remaining brains did not exhibit obvious visible hyperintensities in M1. Representative susceptibility and R2* maps from all brains are shown in Figure S6 in the Supplementary material.

**Figure 7.**
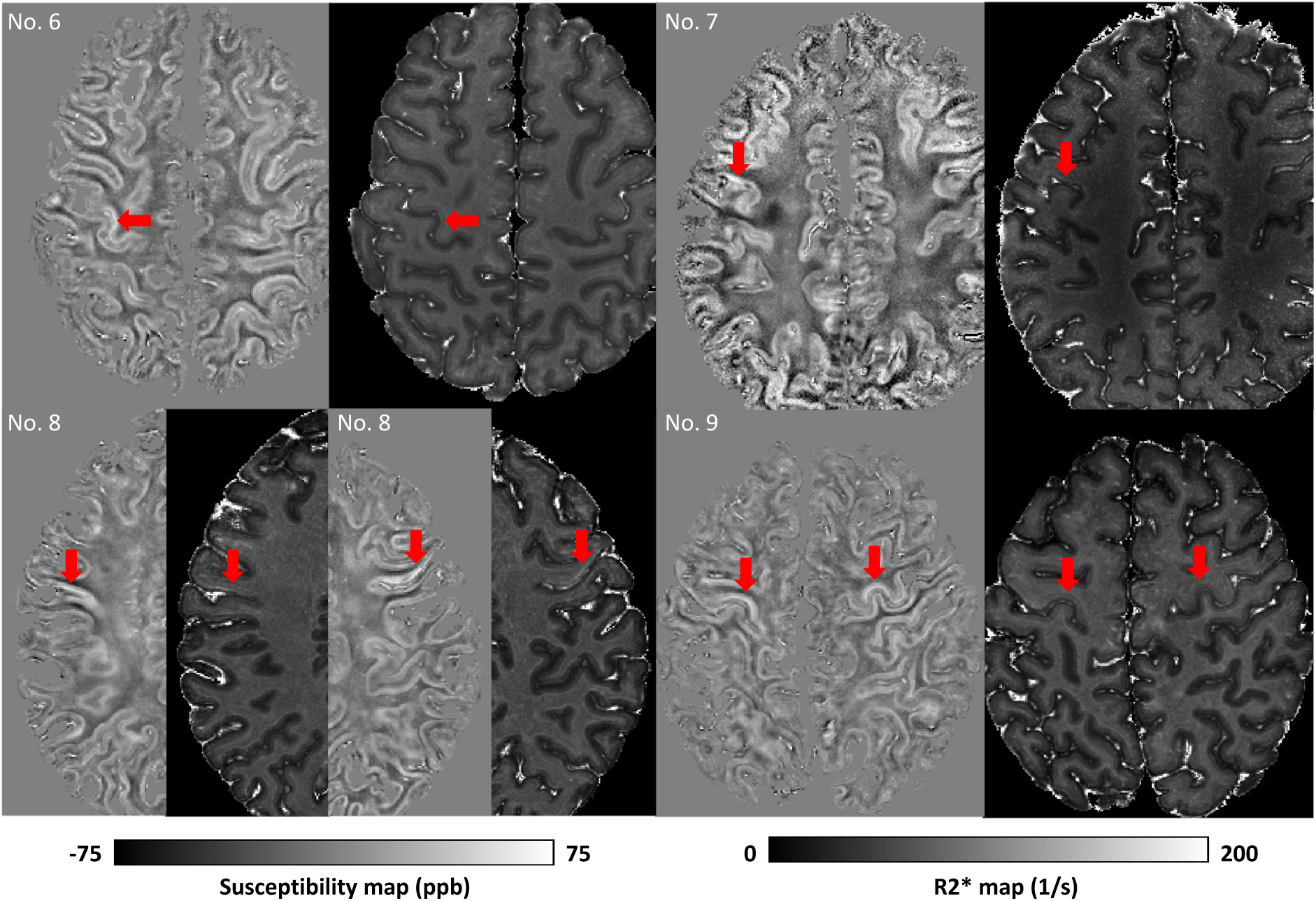
Susceptibility and R2* images from the four ALS brains that exhibited visually apparent hyperintensities in susceptibility and R2* within the cortical grey matter of M1. Red arrows are indicating the cortical hyperintensities. Specifically, hyperintensities were found in the left M1 hand area in brain No. 6, left M1 face area in brain No. 7, both right and left M1 face area in brain No. 8, and both right and left M1 hand areas in brain No. 9.

ALS brains were found to have greater mean susceptibility and R2* values in M1 when compared to controls (Figure 8).

**Figure 8.**
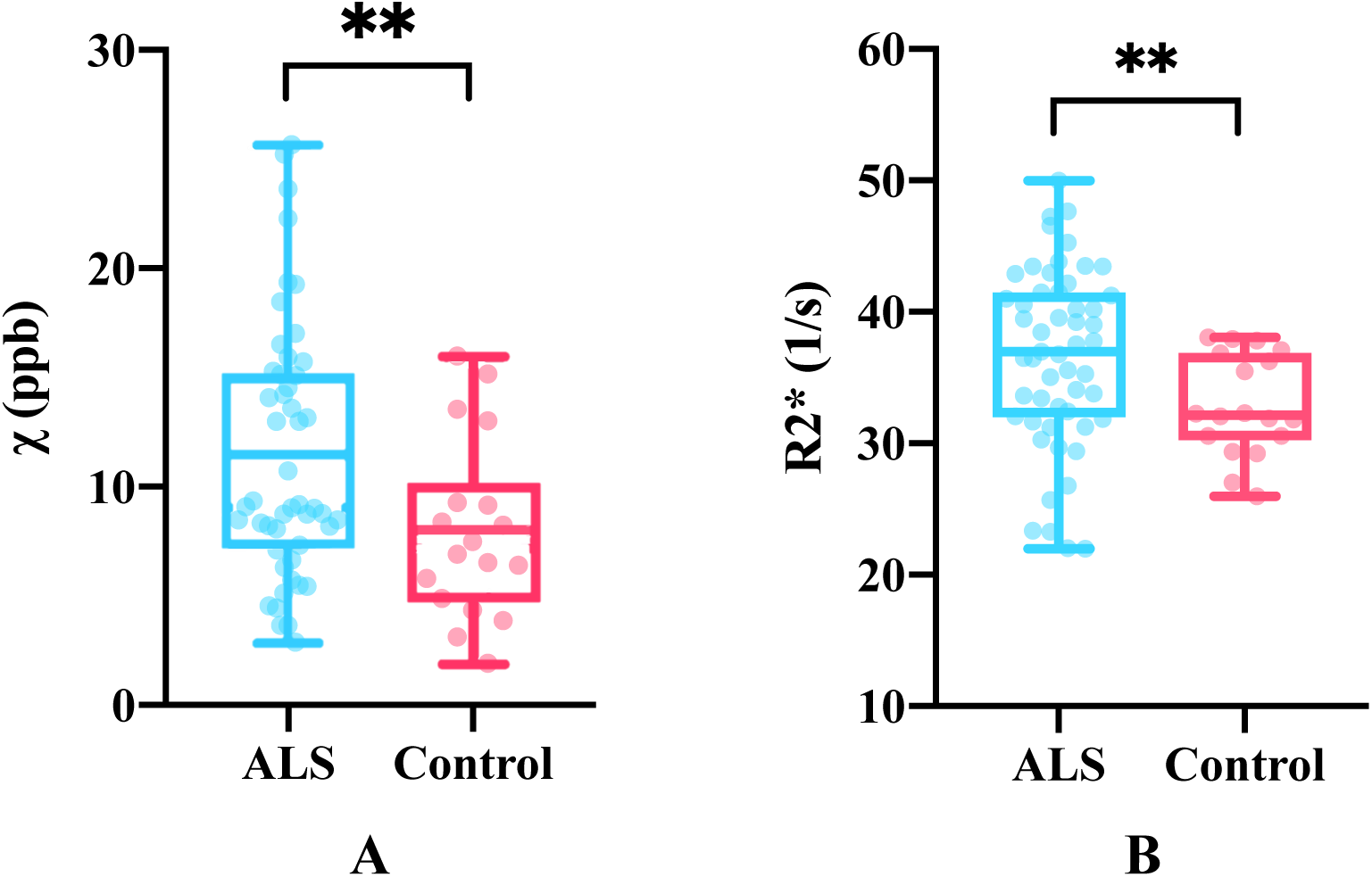
Mean susceptibility (A) and R2* values (B) in all M1 sub-areas in both hemispheres between ALS and control brains. Group differences were significant, with p=0.0098 (Hedges’s g=0.63) and p=0.0065 (Hedges’s g=0.59) for susceptibility and R2* respectively. ** *p<0.01*

The mean susceptibility and R2* values were compared across all regions (M1, ACC and V2) and showed a strong positive correlation (Figure 9). This consistency is expected and reflects the common factors driving both QSM and R2*, which relate to relative susceptibility offsets in different tissue compartments.

**Figure 9.**
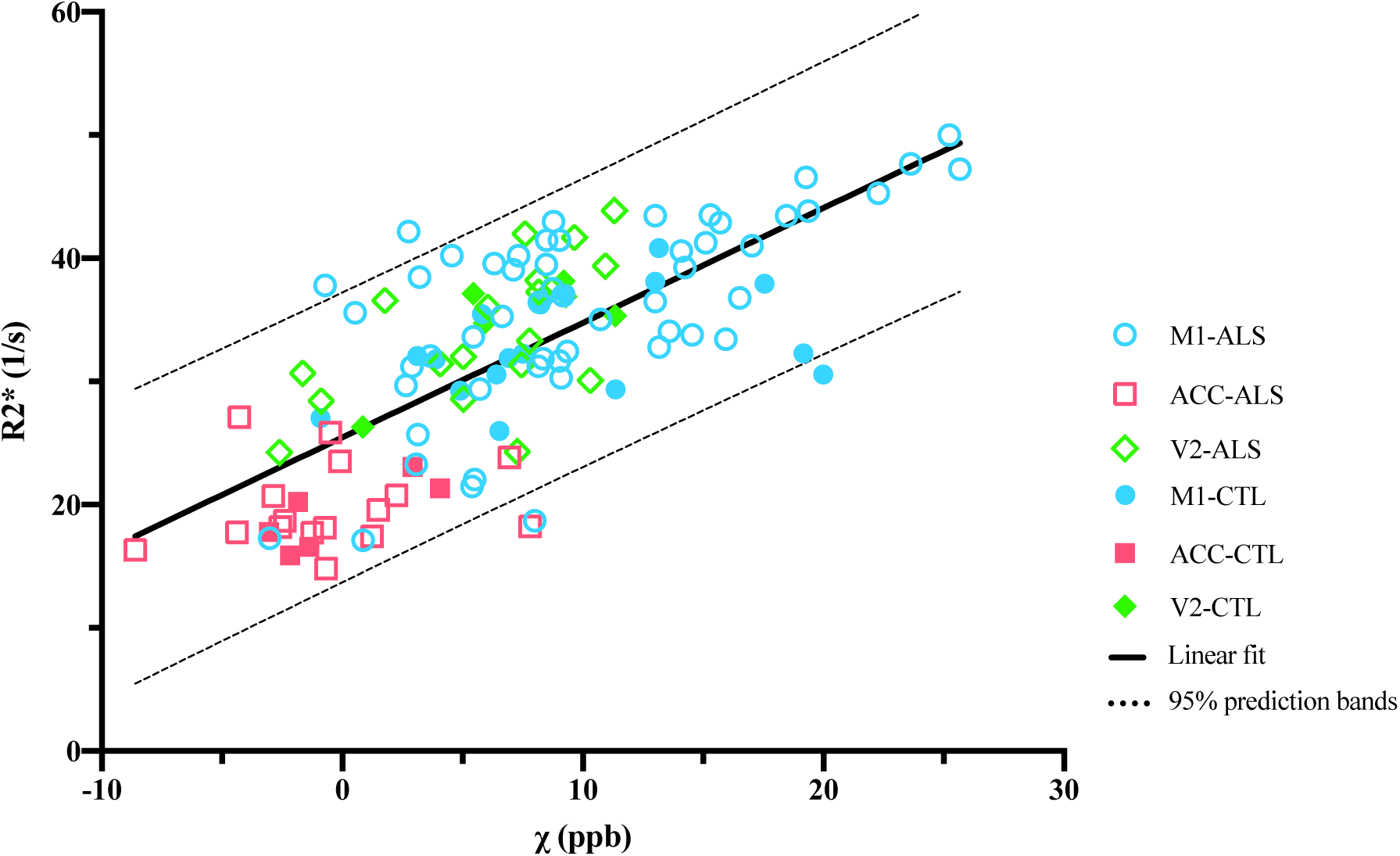
Correlation between mean R2* and susceptibility values within all the ROIs (R=0.73, p<0.0001).

### 3.2 Histological results and their relation to MRI-derived measures

As shown in Figure 10, the cortical grey matter of M1 in ALS brains exhibited higher average ferritin and PLP stained area fractions compared to control brains, although the group differences were not significant. Both susceptibility and R2* showed positive correlations with ferritin and PLP, when the leg and face area of M1, ACC and V2 regions were analysed together (Figure 11 and 12). As both ferritin and PLP contribute to estimated χ and R2* values, and a positive correlation between ferritin and PLP SAFs was found (r=0.35, p=0.011), partial correlation (PLP was regressed out when correlating χ/R2* with ferritin, and vice versa) results are also reported.

**Figure 10.**
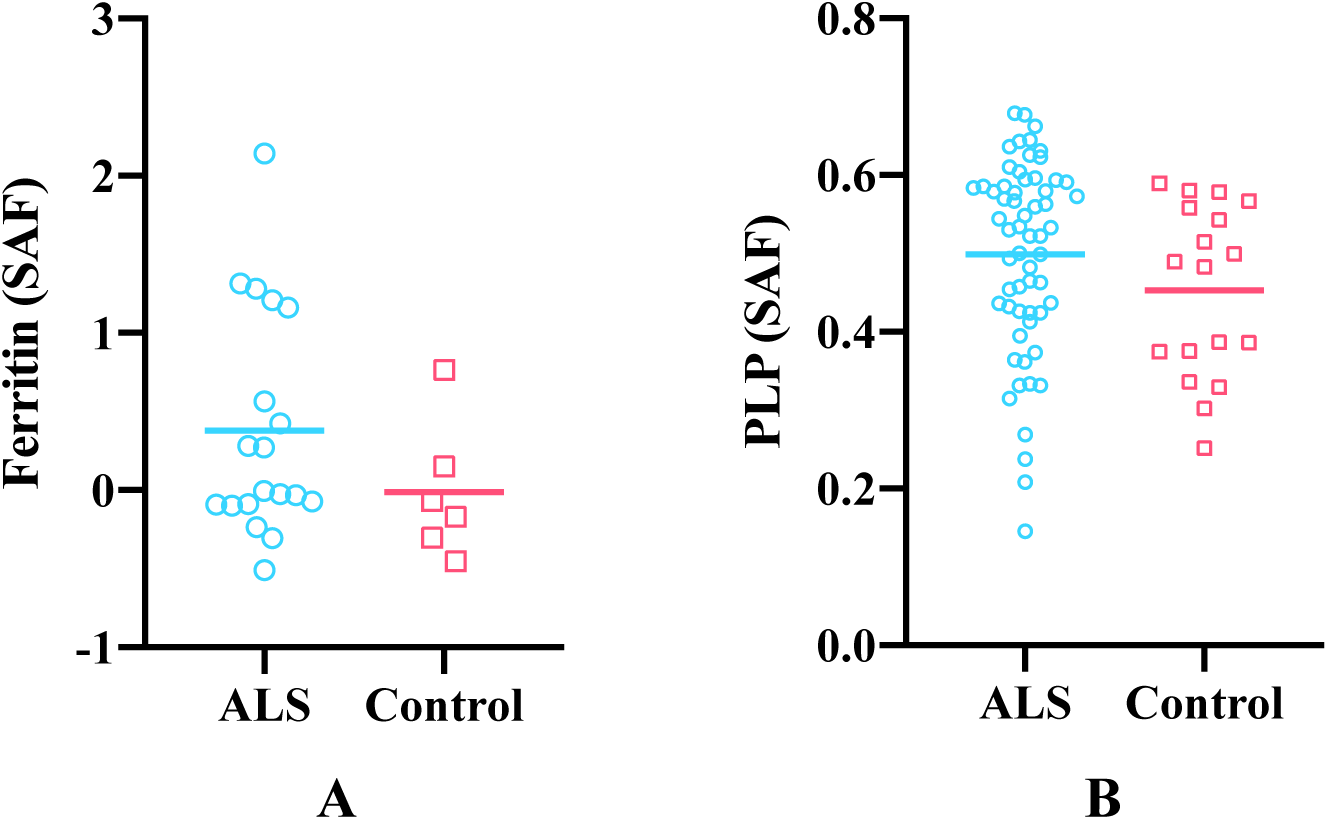
Group mean differences in ferritin (A) and PLP (B) stained area fractions (SAF) between ALS and control brains in M1. Ferritin results are drawn from M1 face and leg areas in the left hemisphere, while PLP results consists of all M1 sub-areas in both hemispheres (reflecting the more manually-intensive nature of the ferritin stains). ALS brains exhibited greater ferritin and PLP SAFs compared to control brains but group differences for both stains were not significant (p=0.125 and p=0.135, respectively).

**Figure 11.**
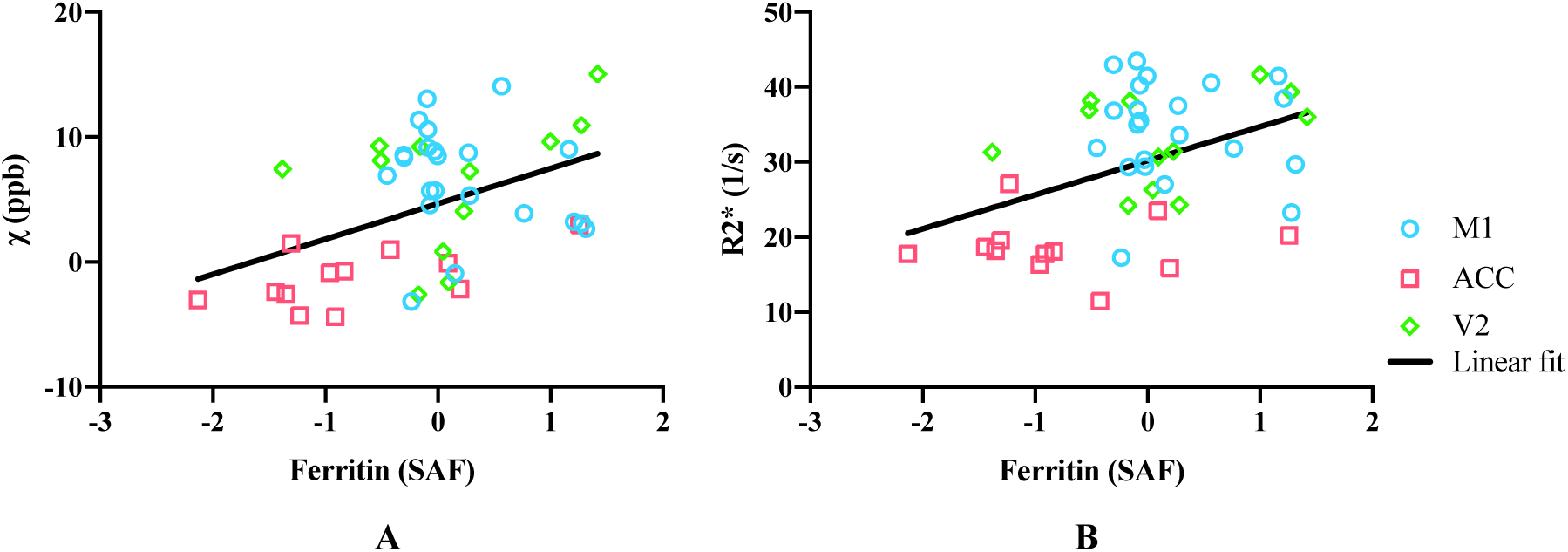
(A) Susceptibility values correlate with ferritin (r=0.43, p=0.0027), including after partial correlation controlling for PLP (r=0.34, p=0.023) when analysing over the three ROIs. When analysing individually, correlation between susceptibility and ferritin was only found in the ACC (r=0.59, p=0.043), no significant correlation was found in M1 or V2 (r=-0.22, p=0.32 and r=0.14, p=0.67, respectively). (B) R2* values correlate with ferritin (r=0.42, p=0.0035), including after partial correlation controlling for PLP (r=0.31, p=0.039) when analysing over the three ROIs. When analysing individually, no significant correlation between R2* and ferritin was found in M1, ACC or V2 (r=-0.056, p=0.80; r=-0.0031, p=0.99 and r=0.30, p=0.35, respectively). M1 includes face and leg areas. Data were from the left hemisphere. Note that Ferritin SAF values were standardised, hence have negative values.

**Figure 12.**
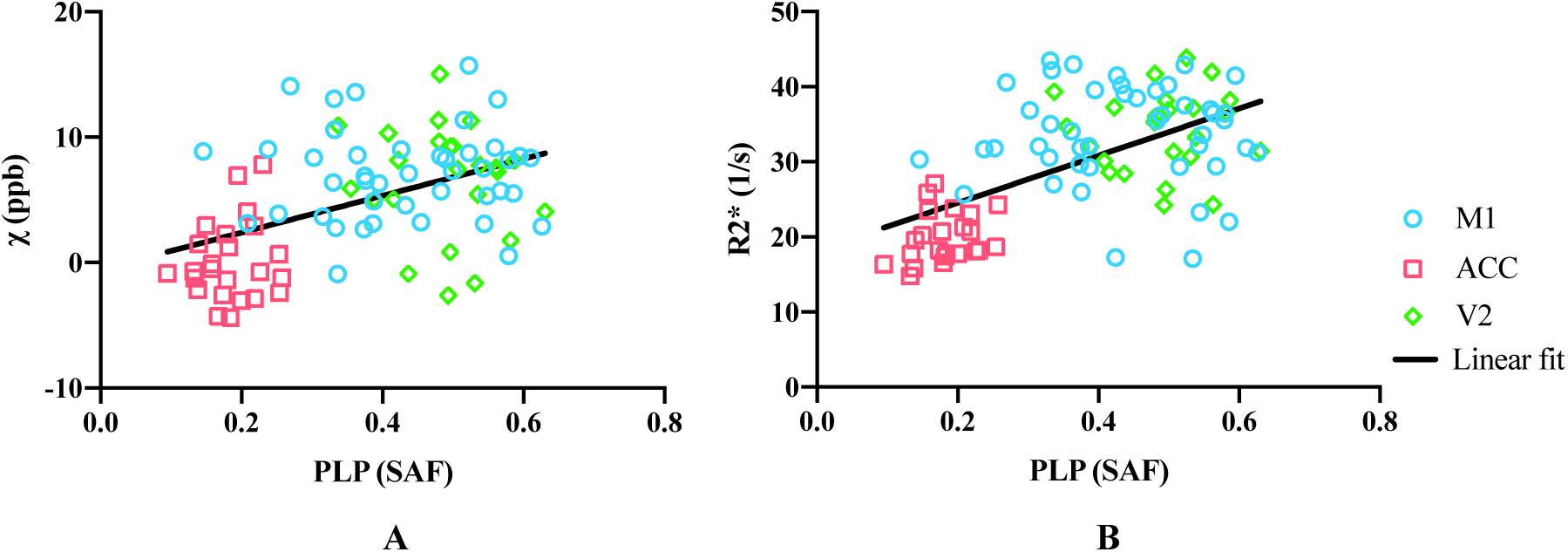
(A) Susceptibility values correlate with PLP (r=0.47, p<0.0001), including after partial correlation in the left hemisphere controlling for ferritin (r=0.37, p=0.015) when analysing over the three ROIs. When analysing individually, no significant correlation between susceptibility and PLP was found in M1, ACC or V2 (r=0.0076, p=0.96; r=0.14, p=0.50 and r=-0.16, p=0.47, respectively). (B) R2* values correlate with PLP (r=0.40, p<0.0001), including after partial correlation in the left hemisphere controlling for ferritin (r=0.48, p=0.00097) when analysing over the three ROIs. When analysing individually, no significant correlation between R2* and PLP was found in M1, ACC or V2 (r=-0.0041, p=0.98; r=0.26, p=0.22 and r=0.039, p=0.86, respectively). M1 includes face and leg areas. Data presented here are from both hemispheres.

## 4 Discussion

Acquisition of high-quality quantitative susceptibility mapping in whole, post-mortem brains has both challenges and opportunities that are not present in vivo. Opportunities include long scan times to increase SNR and spatial resolution, as well as the ability to compare against histology drawn from a broad range of brain regions. The proposed experimental setup and image processing pipeline for quantitative susceptibility and R2* mapping was developed to address the specific challenges associated with post-mortem acquisition. First, the absence of skull will in general alter the macroscopic background field compared to in vivo. By embedding the brain within a large sphere, we can improve the overall shim quality. Second, reliable positioning of the brain is important for achieving consistent QSM results, particularly in the white matter where susceptibility anisotropy causes the estimated susceptibility value to depend on orientation to the B0 field. This challenge was overcome through the use of an internal brain-shaped holder that could be reliably positioned within the outer sphere. Third, the presence of air bubbles causes severe signal dropout and phase aliasing, while small air bubbles induce strong dipole-shaped fields around the region that may lead to streaking artefacts during the dipole inversion step. Finally, although true head motion is not an issue post-mortem, the long scan times lead to image misalignment due to B0 drift and linear phase shifts. This has been addressed by appropriately constrained post-processing corrections.

Similarly, histology data requires optimisation. For both ferritin and PLP, non-specific (background) DAB staining was avoided using treatment with 3% H2O2 to quench peroxidase activity, which can interact with DAB. The quality of immunohistochemistry was assessed during protocol development by an experienced histologist, and validated by a clinical neuropathologist. We now describe for each stain the cellular specificity that was observed for DAB and prior literature reporting similar results. PLP staining was observed to be specific to axonal fibres of the cortical grey matter and no staining was observed in neurons or oligodendrocytes. Myelin PLP staining was most dense in M1 and V2 and less dense in ACC. This agrees with previous reports that ACC is substantially less myelinated than the M1 and V2 [46, 47]. In PLP, the majority of the axonal fibres were observed to run in vertical and horizontal patterns with some bundling of vertical fibres, consistent with previous findings [48]. Ferritin staining was apparent in myelin, glial cells and oligodendrocytes. Previous work showed that myelinated fibres colocalised with ferritin staining in both grey and white matter [49]. Furthermore, microglial ferritin staining has been demonstrated in the frontal cortex of Alzheimer’s disease [50] and in the deep layers of the motor cortex of ALS [24]. Since the majority of iron is bound to ferritin, studies have validated intracellular ferritin as a surrogate marker for iron content in tissue [50, 51]. Shortcomings of immunostaining with respect to quantification are described below.

Susceptibility and R2* both showed positive correlation with the ferritin stained area fraction (Figure 11). This is as expected, due to the paramagnetic properties of iron [13, 14], what was not expected, however, is that both susceptibility and R2* values additionally showed positive correlations with PLP. Positive correlation is expected for R2* estimates [52], but a negative correlation would be predicted for susceptibility due to the diamagnetic properties of myelin. A similar result to ours has been previously reported [20], which has been attributed to a correlation between iron and myelin content in grey matter, which was also observed in our data. To explore whether this explanation is consistent with our data (i.e. that the apparent correlation was in effect an indirect relationship mediated by cortical iron content), we performed a partial correlation analysis. Regressing the ferritin SAF out of both PLP SAF and susceptibility estimates prior to correlation reduced the positive correlation, but did not produce the expected negative correlation. To the extent that ferritin SAF accounts for iron (caveats on this point are discussed below), the partial correlation analysis was unable to explain this finding. This positive partial correlation of PLP with susceptibility is not in agreement with the diamagnetic nature of myelin and remains unexplained.

More generally, it is worth noting that the correlations presented here are driven primarily by regional differences rather than variation within region, reflected as a clustering of regions in Figure 11. In this situation, correlations are prone to identifying apparent relationships between any properties exhibiting regionally-specific differences. In general, we did not observe meaningful correlations within individual regions, with the only significant correlation being between ferritin SAFs and susceptibility in the ACC, although this does agree well with the overall inter-regional correlations. The lack of significant correlations in individual regions could reflect low statistical power in the presence of small effect size (differences between brains) and small samples (n=12).

This unexpected correlation between PLP SAF and susceptibility could reflect some limitations in the immunohistochemistry methods used within our study. First, DAB staining does not follow the Beer-Lambert law (the optical density is not representative of the protein concentration). This is the primary motivation for semi-quantitative analysis based on SAF rather than optical density. One drawback of measuring SAF is the need to use thresholding to determine positively stained area. In this work, a single threshold for each stain was manually chosen by a histologist which could potentially fail to account for slide to slide variability. A pipeline that enables automated thresholding of digital histology images is currently under development [53, 54]. Second, ferritin staining is not directly quantitative of iron load. Each ferritin complex can carry a variable number of iron atoms, meaning that ferritin-associated iron is not necessarily proportionate to the protein content measured here. Third, although ferritin is the principal iron storage protein, non-ferritin iron would also contribute to MR susceptibility signal. This would lead to both an underestimate in the apparent correlation with iron load and, potentially, an imperfect deconfounding of iron from PLP stains. Other techniques that more directly measure iron concentration, such as the mass spectrometry technique used in Langkammer et al. [13], could offer a more accurate quantification of iron in tissue. However, such techniques require heating the sample which may damage the tissue chemical structure and preventing further use of the histology sample.

Previous studies have reported susceptibility and R2* hyperintensities in M1 of ALS patients [25-27]. We observed similar effects in cortical grey matter of M1 in four out of nine ALS brains, corresponding to different sub-regions. This variability might relate to disease heterogeneity. Furthermore, in our study, ALS brains were found to have greater mean susceptibility and R2* values in M1 compared with control brains, while ALS brains also exhibited greater ferritin and PLP stained area fractions compared with control brains, although group differences were not statistically significant. The result of higher PLP SAFs in the M1 region in ALS brains compared to control brains (Figure 10B) is potentially surprising as neurodegeneration is more often associated with demyelination. A related result has been previously reported by Meadowcroft et al. [55], where an increased staining and optical density of subcortical white matter in the primary motor cortex was found in ALS tissue compared to controls using a different myelin stain (Luxol fast blue). Disease-related iron and myelin changes in M1 and their contributions to the MRI susceptibility signal need further investigation.

This study had several limitations. First, the GRE acquisition protocol could be better optimised, as the readout train included unnecessary dead time. To give a feel for the potential gains, if one wished to leave the exact TEs and TR unchanged (e.g. if the first echo required a particularly short TE=2ms), a bandwidth of 210 Hz/pixel could be achieved for echoes 2-5, increasing the SNR by 1.76x for these echoes. Second, registration between histology slides and MRI data was not performed, requiring us to correlate at the level of region-of-interest rather than pixel-wise. A pipeline that enables automated registration of histology slides to 3D MRI images using dissection photos as an intermediary is currently under development [56]. This registration pipeline aims to enable pixel-wise comparisons between MRI and histology acquired within the same sample. Third, we were only able to obtain ferritin histology in limited regions of interest over two separate experiments. This limitation reflects challenges in ferritin staining, with quality of stain being operator dependent such that direct comparison requires that all slides are processed as a single batch. To enable combination of stained area fractions measured from two batches, ferritin SAFs were standardized statistically within batch to account for technician-related bias. Fourth, QSM is a measure of tissue’s relative susceptibility, which is generally made in comparison to an internal reference region (e.g. cerebrospinal fluid). In our study, we lack a reliable reference region that can be confidently considered to be free of pathology (indeed, even the region chosen to represent very low pathology, V2, may be affected). As such, susceptibility values were referenced to the whole brain by setting the mean susceptibility of whole brain to zero. Fifth, only three control brains are included, which limits statistical power in this current cohort. Sixth, we observed a variation of R2* values with distance to the tissue surface in post-mortem brains (Figure S7 in the Supplementary material). This may be due to the time it takes for fixative to penetrate into the tissue [57] combined with leakage of the fixative from the tissue surface [58]. We are developing a regression-based correction for this effect based on forward simulation of the diffusion of fixative over time.

## 5 Conclusion

We have developed a QSM and R2* processing pipeline for whole, fixed post-mortem brain acquisition. Comparison with semi-quantitative ferritin and myelin histology demonstrates the potential for susceptibility and R2* maps to serve as a quantitative imaging marker for the diagnosis of ALS or other neurodegenerative diseases.

## Acknowledgements

This study was funded by a Wellcome Trust Senior Research Fellowship (202788/Z/16/Z) and a Medical Research Council grant (MR/K02213X/1). Brain samples were provided by the Oxford Brain Bank (BBN004.29852). The Wellcome Centre for Integrative Neuroimaging is supported by core funding from the Wellcome Trust (203139/Z/16/Z). C. W. is funded in part by the China Scholarship Council (CSC). M.C. is funded by the Royal Academy of Engineering (RF201617\16\23).

We are grateful to Dr Alard Roebroeck for providing the 3D printed shell. We acknowledge the Oxford Brain Bank, supported by the Medical Research Council (MRC), Brains for Dementia Research (BDR) (Alzheimer Society and Alzheimer Research UK), and the NIHR Oxford Biomedical Research Centre. The views expressed are those of the authors and not necessarily those of the NHS, the NIHR or the Department of Health.

