## Supplementary Material for "Methods for quantitative susceptibility and R2* mapping in whole post-mortem brains at 7T"

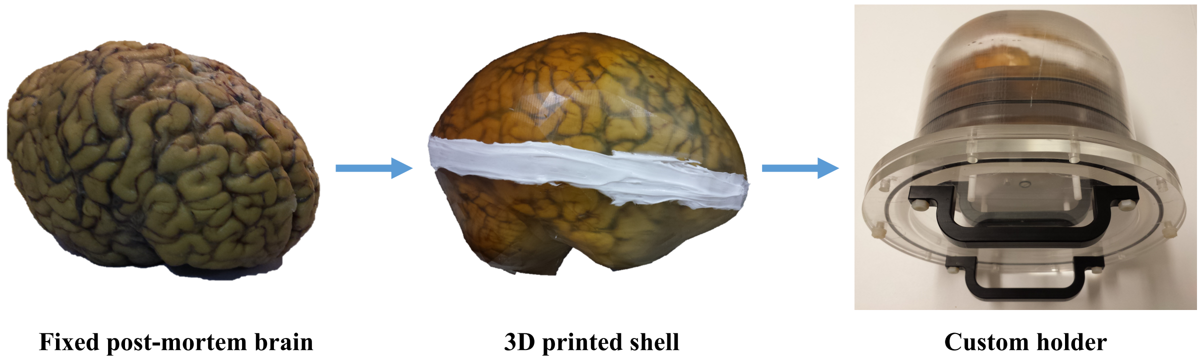


**Figure S1** Illustration of the 3D printed shell and custom holder for scan preparation. The 3D printed shell was provided by Dr Alard Roebroeck, Maastricht University.

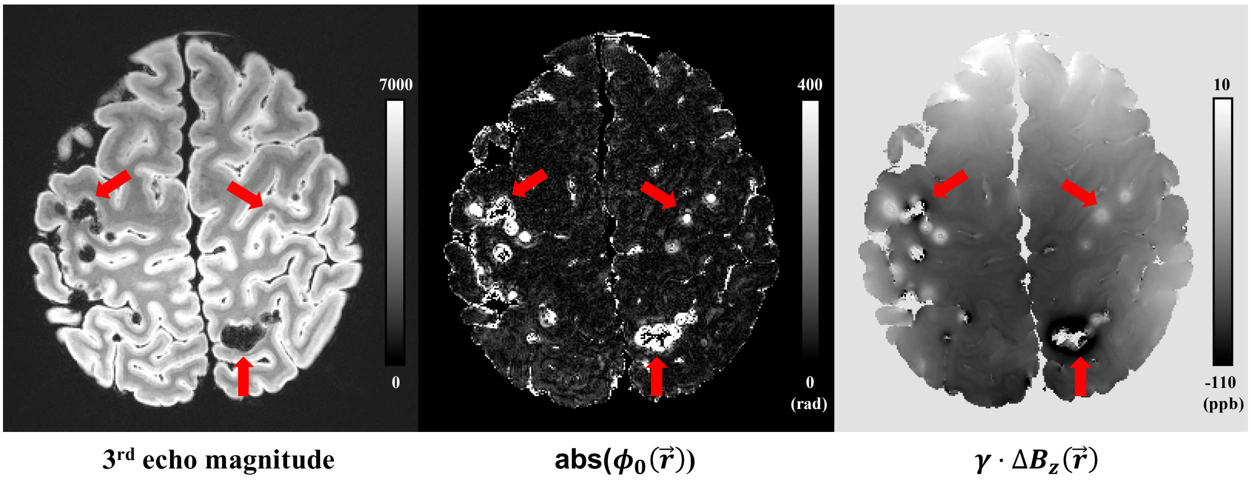


**Figure S2** Illustration of detecting noisy voxels (with unreliable phase that correspond to air bubbles or blood vessels) from the $\phi_0(\vec{r})$ map estimated from fitting. Air bubbles and blood vessels correspond to large, focal voxels on the $abs[\phi_0(\vec{r})]$ maps (red arrows).

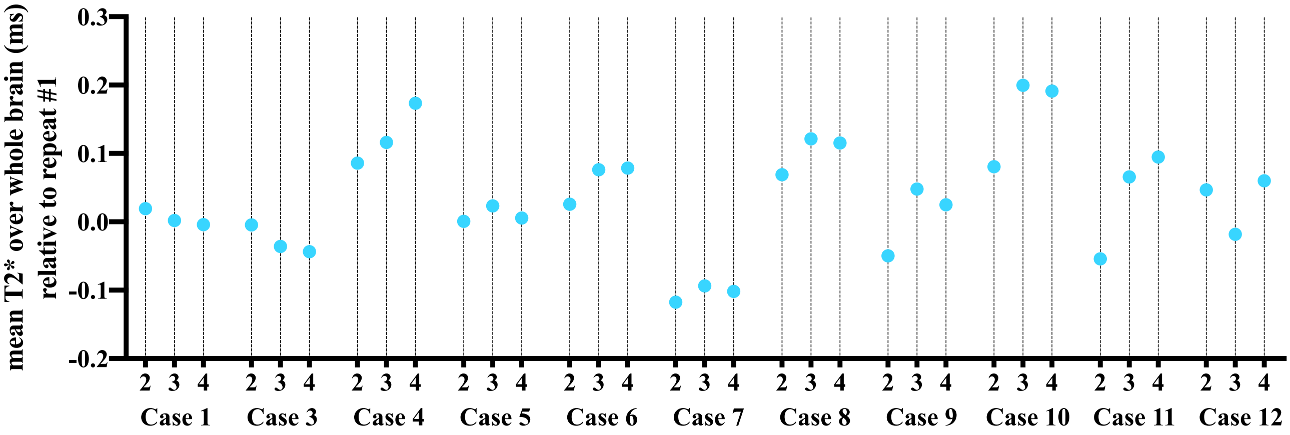


**Figure S3** Change in T2* over the whole brain for repeats #2-4 relative to repeat #1 in 11 brains. Only two repeats were available for case 2, which was not included here.

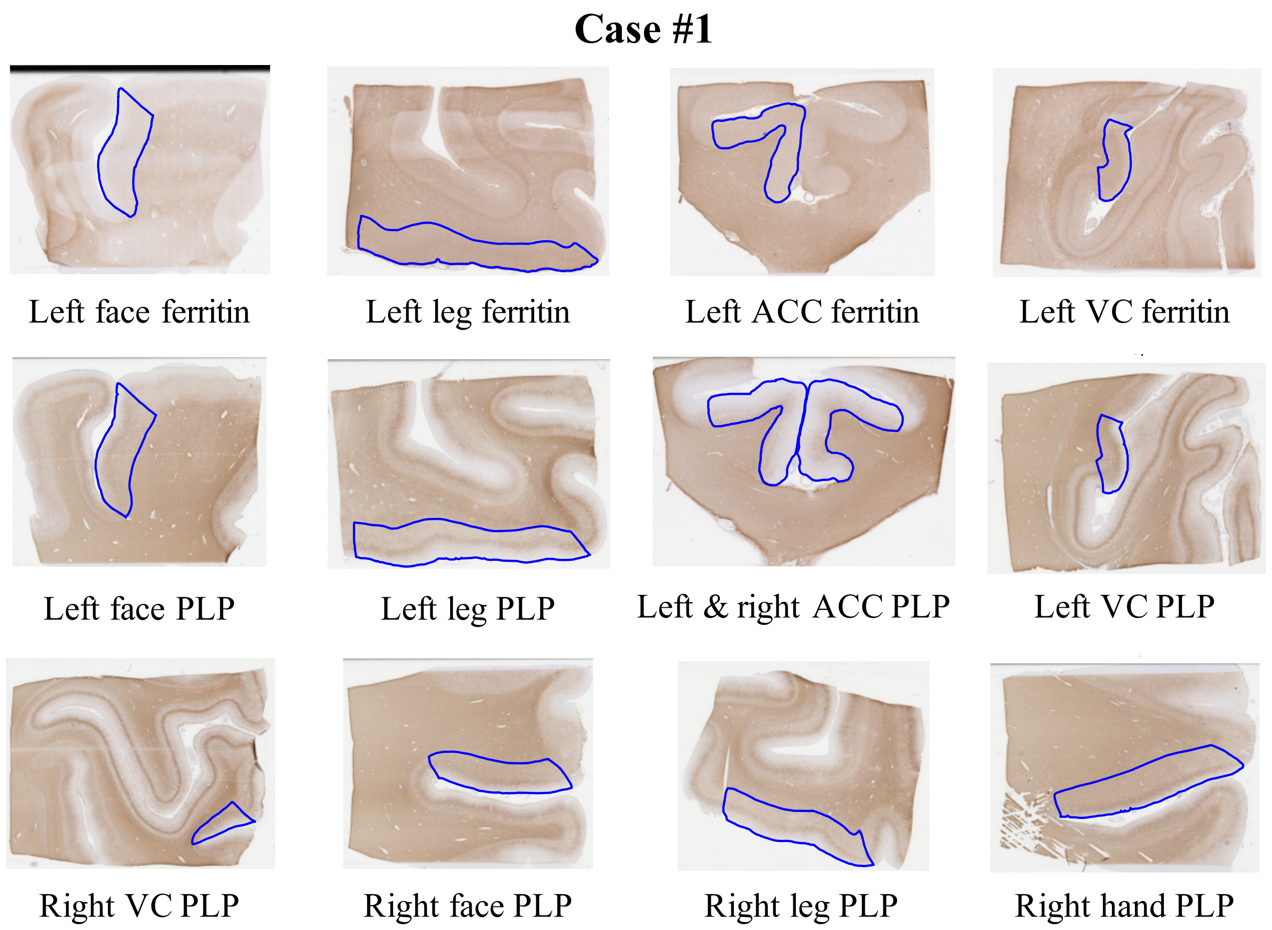


**Figure S4** Digital stained histology images with cortical annotations for Case #1.

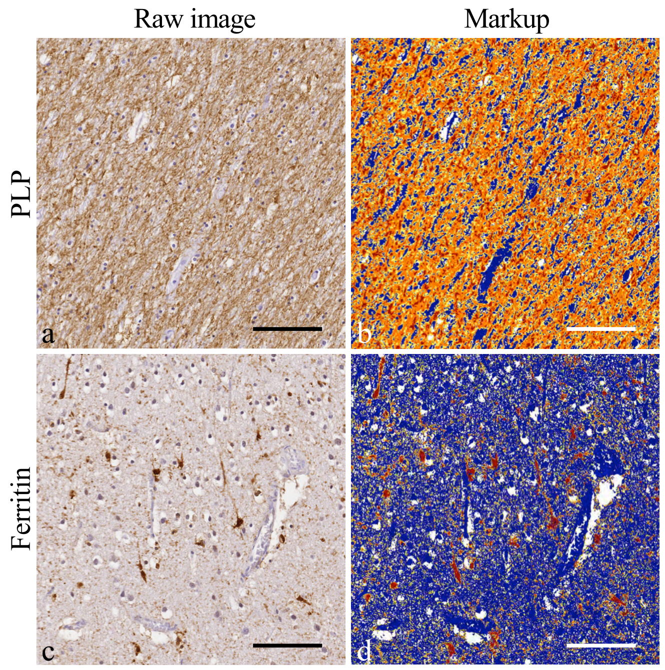


**Figure S5** Digital histology images represent the motor cortex subcortical white matter stained for PLP (a) and grey matter stained for ferritin (c) in ALS. Colour deconvolution for positive staining (brown) generates markup images (b and d) showing unstained pixels as white, negative stained pixels as blue, weak positive pixels as yellow, medium positive pixels as orange and strong positive pixels as red.

*Scale bar = 100 μm*

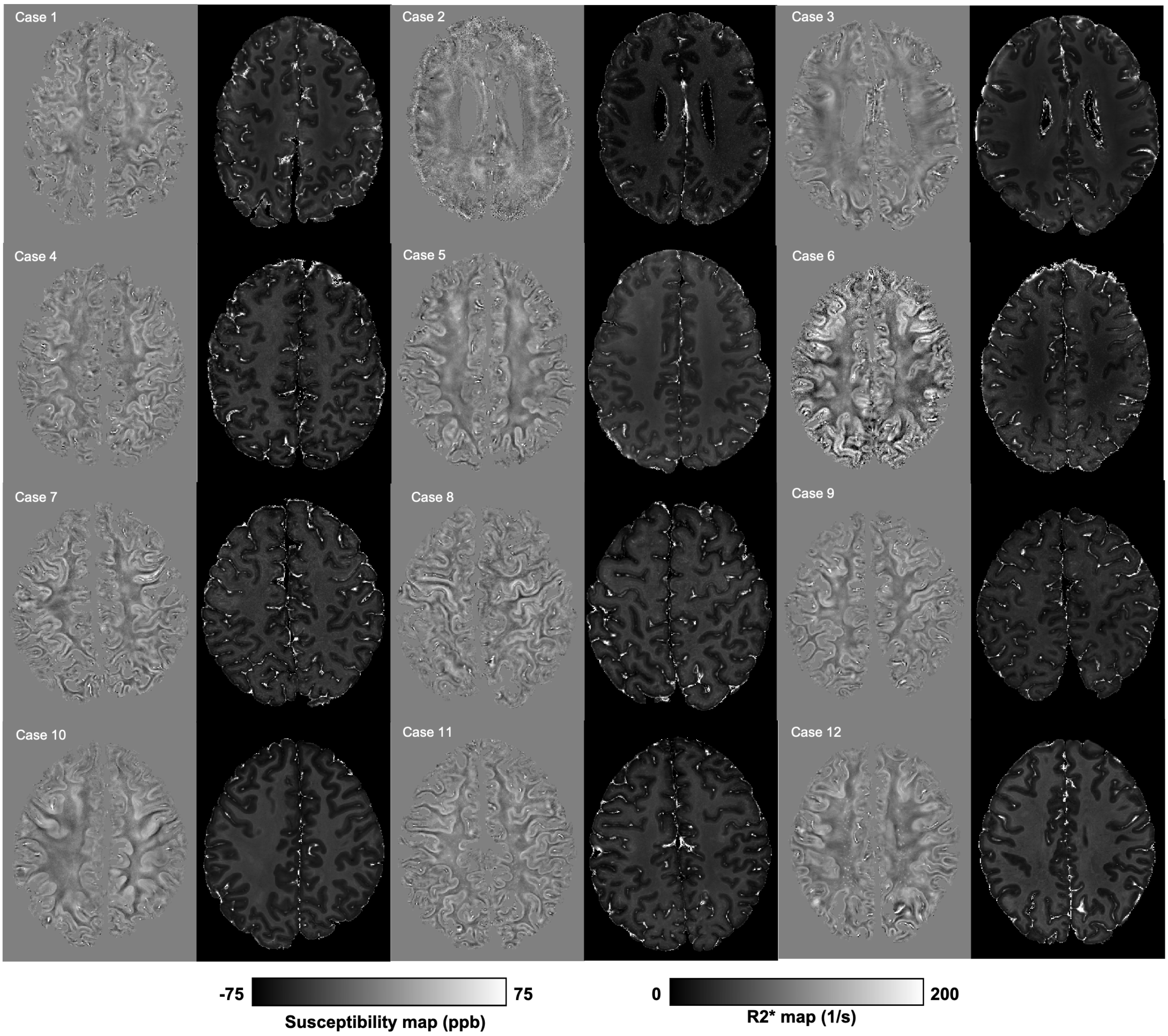


**Figure S6** R2* and quantitative susceptibility maps from all brains.

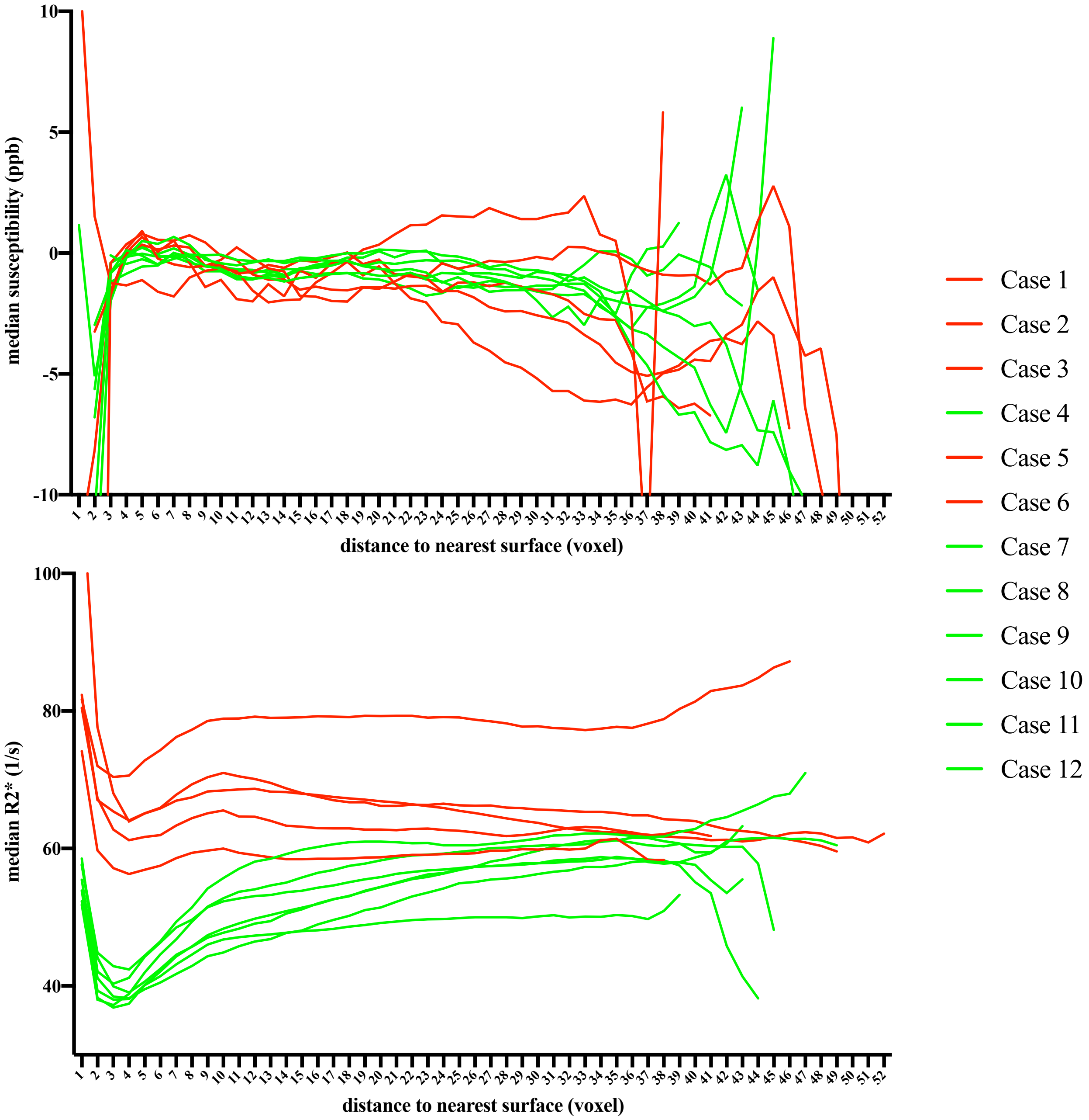


**Figure S7** Median susceptibility and R2* of voxels with different distances to the nearest brain surface for all thirteen brains. Here the red and green lines correspond to the two different fixatives used within our project (red - 10% formalin and green - 10% neutral buffered formalin). For the R2* maps, a difference is observed for the R2* estimates which depends on the type of fixative used. A similar trend is not observed for the susceptibility estimates.
